# The Role of the Public Health Service Equalization Program in the Control of Hypertension in China: Results from a Cross-sectional Health Interview Survey

**DOI:** 10.1101/631796

**Authors:** Jiangmei Qin, Yanchun Zhang, Masha Fridman, Kim Sweeny, Lifang Zhang, Lu Mao, Chunmei Lin

## Abstract

**Objectives:** Non-communicable diseases (NCDs) have become the main cause of mortality in China. In 2009, the Chinese government introduced the Public Health Service Equalization (PHSE) program to restore the primary healthcare system in both essential medical care and public health service provision. This study evaluates the impact of management on hypertension control and evaluate how the program works.

**Methods:** The China National Health Development Research Centre (CNHDRC) undertook the Cross-sectional Health Service Interview Survey (CHSIS) of 62,097 people from primary healthcare reform pilot areas, across 17 provinces from eastern, central and western parts of China in 2014. This study is based on CHSIS survey responses from 9,607 participants, who had been diagnosed with hypertension. Regression analysis was used to estimate the impact of management provided under PHSE on hypertension control adjusting for the effects of other known determinants of hypertension control.

**Findings:** Uncontrolled hypertension was markedly lower among respondents, whose hypertension had been managed (22.4% in managed patients versus 31.1% in unmanaged patients, p<0.001). The interaction between PHSE management and the geographical region was highly significant in the model (p<0.001), suggesting that the PHSE program was not equally effective in all regions. Further analysis suggested that approximately 10% of regional variability was attributed to differences in administrative systems, as there was a significant association (P=0.014) between the presence of established regional Information Management Systems (IMS) and increased PHSE effectiveness. Insurance (χ^2^(5)=4.4, p=0.496) and Hukou (χ^2^(1)=2.4, p=0.121), which denote social security and urban rural differences, respectively, were not significant predictor of hypertension control.

**Conclusion:** Active management of hypertension through the PHSE program was effective with 7.31 million more patients receiving hypertension control and equalization of service delivery was reflected to some extent. The link between established IMS and regional variability in the impact of PHSE highlights the importance of effective management of patient referrals and follow-up. Further investigation is needed to explore the factors that influence the effectiveness of PHSE.

## 1. Introduction

The leading causes of mortality in China have shifted relatively quickly from infectious diseases and perinatal conditions to chronic diseases and injuries [1,2]. This has been accompanied by an increase in hypertension and other cardiovascular risk factors which were responsible for 1.7 million deaths from stroke and 948,700 deaths from ischemic heart diseases in 2010 [3], making these factors the leading causes of death in China. In recognition of the failure of the prevailing health system at the time, the central government introduced a new healthcare reform plan in 2009 to restore the primary healthcare system in both essential medical care and public health service provision [4]. One important measure was the program entitled “Public Health Service Equalization for All (PHSE)” which supports community health organizations to deliver a defined package of basic health services throughout the country [5]. In urban areas, these organizations are called community health centers and stations; in rural areas they are township health centers and village clinics. This essential health care package was designed to equalize essential health care packages, and focused on maternal and child health, elderly people, and chronic disease patients. A major aim of the program is to combat the increasing burden imposed by non-communicable diseases (NCDs), through a range of measures including health education and management of hypertension and diabetes [6], similar to the recommendations by the WHO for essential packages of interventions for non-communicable diseases by primary care facilities [7].

Funding for the program was provided by the Government initially on the basis of 15 Chinese yuan (CNY) per capita each year, which was increased to 20 CNY in 2011 and 50 CNY in 2017 [8]. From 2009 to 2013, over 140 billion CNY (or around USD 21 billion) was invested in this program. It has been estimated that about 18% of this investment was spent on management of patients with hypertension for a total of around 25.2 billion RMB (or USD 3.8 billion) [9]. By the end of 2013, the number of primary health care facilities providing services under the PHSE reached 2.96 million, including 476,073 community health service centers (or stations), 1,244,054 township hospitals, and 1,238,022 village clinics [10]. The specifications for the PHSE and the requirements for service delivery were revised in 2011, 2013 and 2017, ensuring that hypertensive patients aged 35 and over, be followed-up, monitored and evaluated on a regular basis. However, little is known about the effectiveness and benefit of such a large investment, especially from the population’s perspective.

A longitudinal cohort study of the Chinese population, the China Health and Retirement Longitudinal Study (CHARLES) conducted in 2015 demonstrated that factors such as age, sex, smoking habits, drinking habits, household income, health insurance, BMI, residential region, marital status, educational level and nationality were significantly associated with the status of hypertension: awareness, treatment and control [11]. A recent cross-sectional study [12] discovered that a lower likelihood of awareness and treatment of hypertension was associated with younger age, lower income, males, with an absence of previous cardiovascular events, diabetes, obesity, or alcohol use.

Also the rate of control of hypertension was universally low across all subgroups. Data from the CHARLS has highlighted the importance of health insurance in the ability of patients to control hypertension [13]. However, very few evaluations have focused on the role of primary care facilities in hypertension control, although primary care in low-resource settings is considered to be the backbone for implementation of essential NCD interventions [7,14].

Two related reports have described the progress of the PHSE program. The Annual Report of Essential Public Health Services Performance Evaluation has been conducted every year since 2010 by the Center for Project Supervision and Management of National Health and Family Planning Commission (NHFPC) [15]. It reported that from 2011 to 2013, the number of hypertensive patients managed by the program increased from 65,864 million to 85,030 million. Another report conducted by the Community Health Association of China (CHAC) in 2014 [16], was similar to the NHFPC 2013 evaluation. However, none of these studies evaluated the effectiveness of the PHSE from the perspective of hypertension control. The PHSE program facilitates patients seeing the same doctor regularly for management of hypertension. Evidence from the USA shows that the rates of controlled hypertension were significantly higher among persons who visited the same facility for their health care or saw the same provider [17]. This study aims to identify differences in hypertension control between patients managed by the PHSE program and those not managed by the program by using data from the CHSIS.

Based on the studies cited above and others, 12 factors in three categories are considered as determinants of hypertension control in this study. The first category consists of socio-demographic factors influencing adherence to medication, including age, gender, geographical region, health insurance, financial difficulties, and education level. The second category includes disease-related factors, such as smoking, drinking, exercise, blood glucose levels, and the presence of complications and co-morbidities. Hypertension management, denoting medical support and follow-up by a specialist, is the factor considered as the effect of the PHSE program.

This study measures the impact of the PHSE program by comparing the degree of hypertension control among patients managed by the program and those that are outside the program, using the results of the CHSIS. It assesses the contribution of program management to hypertension control while considering other relevant factors suggested by the literature.

## 2. Methodology

### Source of data and sampling

The CHSIS used in this study was conducted in 2014 by the China National Health Development Research Center (CNHDRC) and was funded by the National Health and Family Planning Commission of the People’s Republic of China (NHFPC) and the National Natural Science Foundation (No. 71303173). The primary objective of the survey was to monitor the implementation of primary care reform in China in chosen pilot areas, to draw experiences to promote the integration of the healthcare system, and to disseminate these experiences across the nation. Of the 32 provinces in China, 17 were chosen to participate in the survey based on their representativeness and willingness to participate. These provinces cover the eastern, central and western parts of China considering the rural-urban divide. The basic administrative unit of the financial and tax system in China is the rural county or urban township, so one rural county and one urban district were chosen from a selected city in each province. From the 17 rural counties and 17 urban districts, 10,865 urban households with 30,924 dwellers and 9,912 rural households with 31,173 dwellers were selected by stratified multi-stage sampling resulting in a total of 20,777 households comprising 62,097 persons. Fig 1 provides more detail on the sampling procedure for the survey.

**Fig 1.**
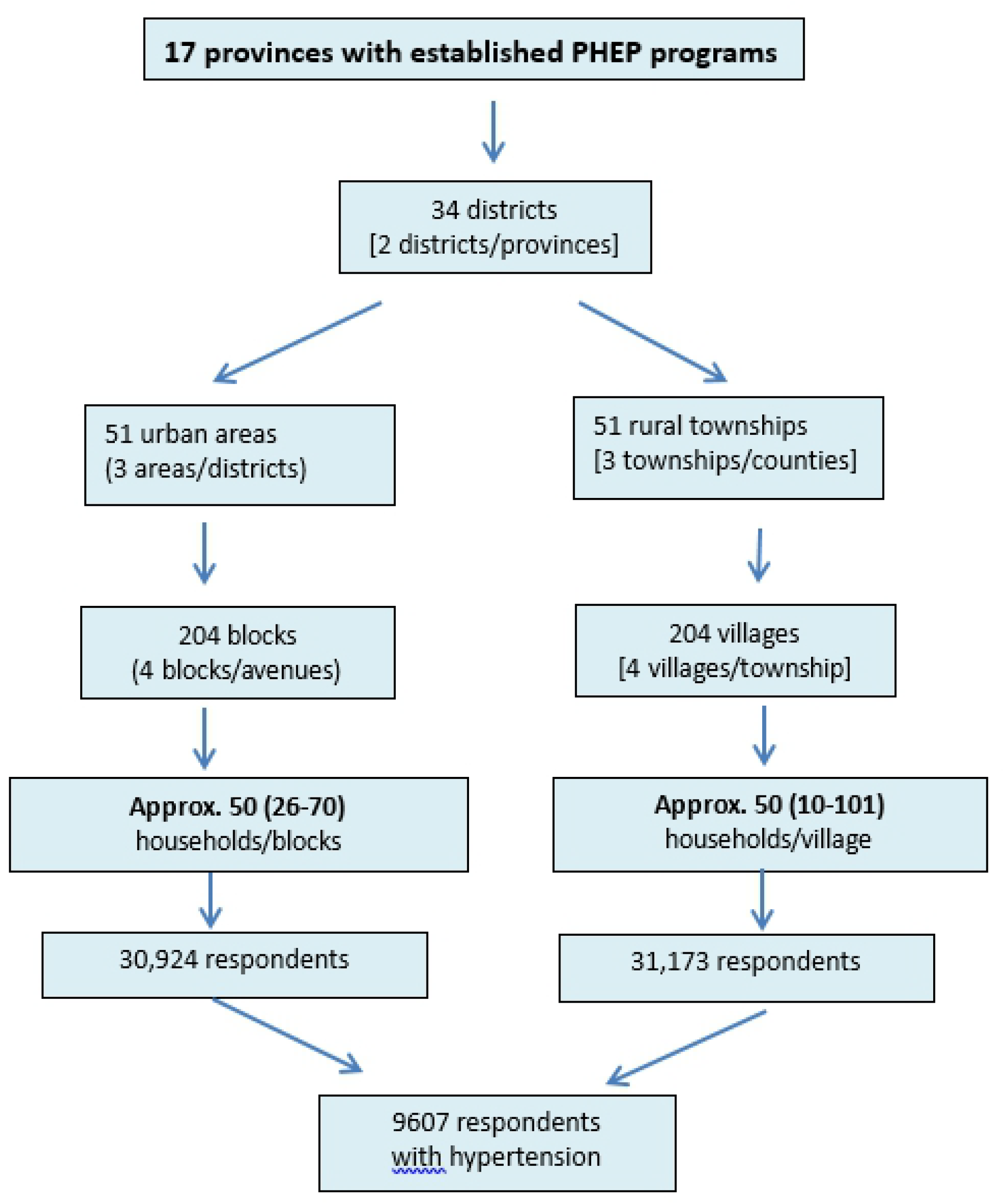
Selection of study participants.

The CHSIS was carried out using face-to-face interviews by undergraduates from medical universities trained by the personnel from CNHDRC. The questionnaire used in the survey was based on the 5th National Health Interview Survey of Households of China conducted in 2013 [18], adjusted to concentrate on primary health care.

In the CHSIS, a total of 9,607 respondents (5155 urban and 4452 rural) aged 15 years and over in the survey answered “Yes” when asked “Have you ever been diagnosed with hypertension by a doctor?”. These respondents form the basis of the analysis in this study.

### Indicators and data method

According to the criteria for public health service delivery developed by the NHFPC for the PHSE, primary health care facilities need to screen people aged 35 years or over for hypertension, and once the person is diagnosed, he or she should be taken into standard management. It requires that patients be seen regularly by a community health care professional at least four times a year, to have their blood pressure measured, to be checked for any signs of deterioration in risk factors, and to obtain advice on behavior and medication. The question “Have any primary health care workers provided advice for your hypertension control during the past three months?” was specifically designed to evaluate hypertension management under the PHSE program. If the answer was “yes”, the patient was considered as being part of the disease management program of the PHSE, and if the answer was “no”, the patient was considered as not being under the disease management program of the PHSE.

Patients were regarded as having hypertension controlled, if they answered “Yes” to the question “Was your blood pressure normal when it was last measured?”. They were considered as not controlled if the answer was “no” or “not clear”.

Other questions addressed demographic characteristics (residence, gender, age, number of household participants), socio-economic status (annual disposable income of the household, health insurance, education level), disease-related health behaviors (smoking, drinking and exercise), self-reported wellbeing and compliance with medication. The questions used to measure these factors are listed in Table 1 and with more details provided in the Supplementary Material (S1Table).

**Table 1.** Factors considered in analysis of hypertension control.

| Factor | Question | Values |
| --- | --- | --- |
| Hukou (Residence) | 1 | Hukou registration: urban, rural |
| Gender | 2 | Male, female |
| Age | 3,4 | Under 35, 35-44, 45-54, 55-64, 65 and over |
| Education | 5 | Education level: No education, primary school, junior high school, senior high school, technical school, middle technical school, senior technical school, university or higher |
| Insurance | 6 | Health insurance coverage: urban employees' health scheme, urban residents' health scheme, new cooperative medical scheme, rural and urban residents' health insurance, other health insurance |
| Wellbeing | 7 | Self-assessed on a scale 0-100. |
| Smoking | 8,9 | None or quit, 1-10, 11-20, 21-30, and 30+ cigarettes per day |
| Drinking | 10 | Yes or no |
| Exercise | 11 | Once, 1-2, 3-5, and 6 times or over a week |
| Management | 14 | Managed, not managed |
| Compliance (Medication) | 15 | Compliance with medication: complete compliance, partial compliance and non-compliance |
| Control | 16 | Controlled, uncontrolled hypertension |
| Income | S1 Table | Household annual disposable income: low, lower-middle, middle, upper-middle, upper, and extreme upper |
| Region | S3 Table | Districts grouped into east urban (EU), east rural (ER), middle urban (MU), middle rural (MR), west urban (WU) and west rural (WR) |
| District | S3 Table | 17 urban districts, 17 rural counties |

The ethics committee of the China National Health Development Research Centre reviewed and approved the present study, and the written informed consent was obtained from each participant before data collection.

### Analytical method

A Pearson chi-square (χ^2^) test was used to compare the managed and unmanaged hypertensive patients for each factor. Results were considered significant if P<0.05. As age and wellbeing were continuous variables, a T-test was used in comparing differences with the significance level also set at P<0.05 (Table 2).

**Table 2.** Comparison of characteristics between Management groups.

| Independent variables | Not managed<br>[N=1,906] | Managed<br>[N=7,601] | Test of difference<br>(p-value) |
| --- | --- | --- | --- |
| Compliance |  |  | X2(1)=70.2<sup>a</sup> (<0.001)<sup>a</sup> |
| Every day | 1,489 (78.1%) | 6,576 (85.4%) |  |
| Sometimes | 311 (16.3%) | 914 (11.9%) |  |
| Never | 106 (5.6%) | 211 (2.7%) |  |
| Gender |  |  | X2(1)=4.7 (0.031)<sup>a</sup> |
| Male | 929 (48.7%) | 3,541 (46.0%) |  |
| Female | 977 (51.3%) | 4,160 (54.0%) |  |
| Age |  |  |  |
| Mean ± SD | 62.8 ± 11.3 | 63.5 ± 10.8 | t = 3.1 (0.002) <sup>b</sup> |
| Median [Interquartile range] | 62 [56-71] | 64 [57-71] | X2(1)=10.5 (0.001) <sup>c</sup> |
| Self-reported wellbeing |  |  |  |
| Mean ± SD | 70.1 ± 15.7 | 70.9 ± 15.2 | t = 2.0 (0.046)<sup>b</sup> |
| Median [Interquartile range] | 70 [60-80] | 70 [60-80] | X2(1)=4.2 (0.038)<sup>c</sup> |
| Education level |  |  | X2(7)=85.9 (<0.001)<sup>a</sup> |
| None | 262 (13.7%) | 1,490 (19.4%) |  |
| Primary | 518 (27.2%) | 2,477 (32.2%) |  |
| Junior high | 593 (31.1%) | 2,117 (27.5%) |  |
| Senior high | 257 (13.5%) | 847 (11.0%) |  |
| Technical | 13 (0.7%) | 23 (0.3%) |  |
| Middle technical | 106 (5.6%) | 279 (3.6%) |  |
| Senior technical | 81 (4.3%) | 259 (3.4%) |  |
| University or higher | 76 (4.0%) | 209 (2.7%) |  |
| Income per capita (CNY) |  |  | X2(5)=29.5 (<0.001)<sup>a</sup> |
| 0 - 4,747 | 110 (5.8%) | 590 (7.7%) |  |
| 4,748 - 10,887 | 443 (23.2%) | 2,102 (27.3%) |  |
| 10,888 - 17,631 | 439 (23.0%) | 1,745 (22.7%) |  |
| 17,632 - 26,937 | 499 (26.2%) | 1,709 (22.2%) |  |
| 26,938 - 50,968 | 363 (19.1%) | 1,360 (17.7%) |  |
| >50,968 | 52 (2.7%) | 195 (2.5%) |  |
| Health insurance |  |  | X2(5)=267.1 (<0.001)<sup>a</sup> |
| None | 37 (1.9%) | 51 (0.7%) |  |
| Urban employee basic | 938 (49.2%) | 2,876 (37.4%) |  |
| Urban residents | 289 (15.2%) | 671 (8.7%) |  |
| New cooperative | 512 (26.9%) | 3,175 (41.2%) |  |
| Urban and rural residents | 116 (6.1%) | 866 (11.3%) |  |
| Other | 14 (0.7%) | 62 (0.8%) |  |
| Exercise |  |  | X2(5)=20.2 (<0.001)<sup>a</sup> |
| Never | 841 (44.1%) | 3,788 (49.2%) |  |
| Up to 1 session/week | 82 (4.3%) | 244 (3.2%) |  |
| 1-2 sessions/week | 109 (5.7%) | 415 (5.4%) |  |
| 3-5 sessions/week | 192 (10.1%) | 774 (10.1%) |  |
| ≥6 sessions/week | 682 (35.8%) | 2,480 (32.2%) |  |
| Smoking |  |  | X2(4)=7.8 (0.099)<sup>a</sup> |
| None | 1,473 (77.3%) | 6,081 (79.0%) |  |
| 1-10 per day | 145 (7.6%) | 621 (8.1%) |  |
| 11-20 per day | 225 (11.8%) | 749 (9.7%) |  |
| 21-30 per day | 29 (1.5%) | 125 (1.6%) |  |
| >30 per day | 34 (1.8%) | 125 (1.6%) |  |
| Alcohol |  |  | X2(1)=1.6 (0.212)<sup>a</sup> |
| <3 occasions/week | 1,662 (87.2%) | 6,795 (88.2%) |  |
| ≥3 occasions /week | 244 (12.8%) | 906 (11.8%) |  |
| Hukou |  |  | X2(1)=255.0 (<0.001)<sup>a</sup> |
| Urban | 1,334 (70.0%) | 3,821 (49.6%) |  |
| Rural | 572 (30.0%) | 3,880 (50.4%) |  |
| Number of participants/household |  |  | X2(4)=4.8 (0.313)<sup>a</sup> |
| 1 | 1,261 (66.2%) | 4,938 (64.1%) |  |
| 2 | 314 (16.5%) | 1,284 (16.7%) |  |
| 3 | 10 (0.5%) | 75 (0.8%) |  |
| 4 | 0 (0%) | 2 (<0.1%) |  |
|  |  |  | X2(33)=1,048.5 (<0.001)<sup>a</sup> |
| Regions (district/county, city) |  |  |  |
| 1. Xicheng, Beijing | 126 (6.6%) | 398 (5.2%) |  |
| 2. Yunhe, Cangzhou | 135 (7.1%) | 193 (2.5%) |  |
| 3. Runzhou, Zhenjiang | 68 (3.6%) | 252 (3.3%) |  |
| 4. Changning, Shanghai | 41 (2.2%) | 403 (5.2%) |  |
| 5. Keqiao, Shaoxing | 56 (2.9%) | 333 (4.3%) |  |
| 6. Meilie, Sanming | 69 (3.6%) | 158 (2.1%) |  |
| 7. Yinzhou, Tieling | 64 (3.4%) | 154 (2.0%) |  |
| 8. Lanshan, Linyi | 44 (2.3%) | 132 (1.7%) |  |
| 9. Wuhou, Chengdu | 15 (0.8%) | 273 (3.5%) |  |
| 10. Hongshan, Chifeng | 71 (3.7%) | 289 (3.8%) |  |
| 11. Wudang, Guiyang | 69 (3.6%) | 111 (1.4%) |  |
| 12. Chengbei, Xining | 79 (4.1%) | 146 (1.9%) |  |
| 13. Yijiang, Wuhu | 120 (6.3%) | 160 (2.1%) |  |
| 14. Yuhu, Shaoshan | 36 (1.9%) | 282 (3.7%) |  |
| 15. Yiling, Yichang | 86 (4.5%) | 114 (1.5%) |  |
| 16. Qing Yunpu, Nanchang | 123 (6.5%) | 160 (2.1%) |  |
| 17. Changyi, Jilin | 131 (6.9%) | 104 (1.4%) |  |
| 18. Pudong, Shanghai | 88 (4.6%) | 354 (4.6%) |  |
| 19. Shengzhou, Shaoxing | 9 (0.5%) | 360 (4.7%) |  |
| 20. Huanghua, Cangzhou | 17 (0.9%) | 281 (3.6%) |  |
| 21. Jurong, Zhenjiang | 21 (1.1%) | 335 (4.4%) |  |
| 22. Shaxian, Sanming | 49 (2.6%) | 191 (2.5%) |  |
| 23. Pinggu, Beijing | 94 (4.9%) | 276 (3.6%) |  |
| 24. Xifeng, Tieling | 19 (1.0%) | 174 (2.3%) |  |
| 25. Yinan, Linyi | 7 (0.4%) | 186 (2.4%) |  |
| 26. Xinjin, Chengdu | 12 (0.6%) | 203 (2.6%) |  |
| 27. Kaiyang, Guiyang | 25 (1.3%) | 208 (2.7%) |  |
| 28. Huangzhong, Xining | 31 (1.6%) | 166 (2.2%) |  |
| 29. Keshiketeng, Chifeng | 26 (1.4%) | 328 (4.3%) |  |
| 30. Yidu, Yichang | 43 (2.3%) | 190 (2.5%) |  |
| 31. Fanchang, Wuhu | 48 (2.5%) | 212 (2.8%) |  |
| 32. Panshi, Jilin | 18 (0.9%) | 181 (2.4%) |  |
| 33. Xiangtan, Shaoshan | 47 (2.5%) | 294 (3.8%) |  |
| 34. Xinjian, Nanchang | 19 (1.0%) | 100 (1.3) |  |

<sup>a</sup> Pearson $\chi^2$ test. Degrees of freedom are indicated by the subscript.
<sup>b</sup> T-test.
<sup>c</sup> Kruskal-Wallis equality-of-populations rank test.

To further analyze the impact of the program, a logistic regression analysis model was used to identify the role of factors in hypertension control, with the dependent variable being a binary variable (1=non-control of hypertension). To make the variables suitable for the logistic regression model, most variables from the survey questionnaire were transformed into binary or ordinal form. The binary variables were hukou or residence (urban or rural), gender (female or male), disease management (yes or no), and drinking alcohol (yes or no). The ordinal variables were income, education level, health insurance, medication, smoking, and exercise, with the first group in each variable set as reference. Age and well-being were included in the regression analysis as continuous variables. The analysis was performed using Stata (version 14).

A likelihood ratio test was used to assess the significance of any interaction effects between hypertension management and the other factors influencing hypertension control. Household random effects and city random effects were also checked where there was a significant cluster effect within households, but not significant within each city (province). The quantitative effect of each impact factor was analyzed for hypertension control. The impact of management was estimated using a main effects logistic regression model with an adjustment for correlation within households (household cluster effects). The 95% confidence interval of predicted probability is also shown.

## 3. Results

### Demographic characteristics

Key demographic characteristics of managed and unmanaged survey participants are described in Table 2. The overall level of uncontrolled hypertension was markedly lower in the managed group of survey participants (22.4% versus 31.1%). The pronounced difference in uncontrolled hypertension could not be attributed to population differences between management groups, as both groups were similar with respect to those demographic characteristics which are known to influence hypertension control. There were strong similarities in compliance with medication (85.4% versus 78.1% in managed and unmanaged groups respectively), gender (46.0% versus 48.7%), age (63.5 versus 62.8 years), wellbeing (70.9% versus 70.1%) and regular alcohol intake (11.8% versus 12.8%). Trends were also similar in proportions across categories of education, income, exercise, smoking and the number of participants per household. The statistical differences between management groups that were detected for most of these variables arose not from meaningful disparities, but from the large sample size (N=9,607), which made tests of significance sensitive to relatively minor differences. More marked differences between management groups were observed between categories of hukou, type of medical insurance, and geographical regions.

### Main factors selected (regression model selection)

A more accurate comparison of uncontrolled hypertension levels in managed and unmanaged groups was performed using multivariate regression modelling to adjust for any population imbalances in the key characteristics that were listed in Table 2. The selection of independent variables for inclusion in the regression models is outlined in Table 3.

**Table 3.** Selection of independent variables for inclusion in a logistic regression model of uncontrolled hypertension.

| Type of effect | Independent variables | Sample size | Degrees of freedom | LR test $\chi^2$ [p-value]<sup>a</sup> |
| --- | --- | --- | --- | --- |
| Selection of main effects | 1. Geographical region | 9,607 | 33 | 572.6 (p<0.001) |
|  | 2. Compliance | 9,607 | 2 | 97.4 (p<0.001) |
|  | 3. Management | 9,607 | 1 | 61.2 (p<0.001) |
|  | 4. Wellbeing | 9,607 | 1 | 52.7 (p<0.001) |
|  | 5. Education | 9,607 | 7 | 34.2 (p<0.001) |
|  | 6. Income | 9,607 | 5 | 26.9 (p<0.001) |
|  | 7. Age | 9,607 | 1 | 14.4 (p<0.001) |
|  | 8. Smoking | 9,607 | 4 | 14.6 (p=0.006) |
|  | 9. Exercise | 9,607 | 4 | 13.0 (p=0.011) |
|  | 10. Insurance | 9,607 | 5 | 4.4 (p=0.496) |
|  | 11. Hukou | 9,607 | 1 | 2.4 (p=0.121) |
|  | 12. Gender | 9,607 | 1 | 0.2 (p=0.664) |
|  | 13. Alcohol | 9,607 | 1 | 0.1 (p=0.828) |
| Main effects model (log likelihood = -4579.6) | Significant IVs (nos. 1-9 above) | 9,607 | 59 | 1,434.8 (p<0.001) |
| Management interaction effects | 14. Geographical region | 9,607 | 33 | 78.2 (p<0.001) |
|  | 15. Compliance | 9,607 | 2 | 0.3 (p=0.849) |
|  | 16. Wellbeing | 9,607 | 1 | 0.5 (p=0.480) |
|  | 17. Education | 9,607 | 7 | 8.5 (p=0.291) |
|  | 18. Income | 9,607 | 5 | 5.1 (p=0.408) |
|  | 19. Age | 9,607 | 1 | 1.7 (p=0.194) |
|  | 20. Smoking | 9,607 | 4 | 4.3 (p=0.362) |
|  | 21. Exercise | 9,607 | 4 | 4.0 (p=0.411) |
| Interaction model (log likelihood = -4540.4) | [Management x Region] terms, significant IVs (nos. 1-9 above) | 9,607 | 92 | 1,535.8 (p<0.001) |
| Household random effects<sup>b</sup> (log likelihood = -4328.6) | 22. Household |  | 1 |  |
|  | Number of participants | 9,607 |  | 423.8 (p<0.001) |
|  | Number of households | 7,867 |  |  |

<sup>a</sup> Likelihood ratio test.
<sup>b</sup> Household random effects were tested in a model that included significant independent variables: nos. 1-9 and no. 14.

Management of hypertension was one of three most important predictors of hypertension control (likelihood ratio test: χ^2^(1)=61.2), along with compliance (χ^2^(1)=97.4) and region (χ^2^(33)=572.6). The only detectable interaction effect was between management and geographical regions (χ^2^(33)=78.2, p<0.001), which indicated that the management program was not equally effective in all regions.

Insurance (χ^2^(5)=4.4, p=0.496) and Hukou (χ^2^(1)=2.4, p=0.121), which denote social security and urban rural differences, respectively, were not significant predictor of hypertension control. It may reflect that the EPHS program had played a role in equalization from the two perspective.

Random effects for 7,867 households (1.2 respondents/household on average) were strong and highly significant (χ^2^(1)= 423.8, p<0.001), suggesting a tendency towards similar levels of hypertension control within the same household. No significant clustering by provinces was detected in the data.

### Main effects in hypertension control

The logistic regression results show that the estimated proportion of patients with uncontrolled hypertension was 27.7%, or 8.6 percentage points lower among participants in the management program (22.4% versus 31.0%, Table 4).

**Table 4.** The impact of Management on non-control risk of hypertension.

| Independent variables | Odds ratio | Predicted proportions | Change in proportions |
| --- | --- | --- | --- |
|  | [95% CI]<sup>a</sup> | [95% CI]<sup>a</sup> | (p-value) |
| Management |  |  |  |
| Not managed | Reference | 31.0% [28.4-35.6] | Reference |
| Managed | 0.59 (<0.001) | 22.4% [21.3-23.5] | -8.6% (p<0.001) |
| Compliance |  |  |  |
| Every day | Reference | 22.2% [21.2-23.1] | Reference |
| Sometimes | 1.63 (<0.001) | 30.3% [27.7-32.8] | 8.1% (p<0.001) |
| Never | 3.00 (<0.001) | 42.0% [36.5-47.5] | 19.8% (p<0.001) |
| Self-reported wellbeing |  |  |  |
| Change/10% in reported wellbeing | 0.88 (<0.001) | Trend shown in Fig 2 | -2.0% (p<0.001) |
| Educational attainment |  |  |  |
| None | Reference | 27.4% [25.2-29.5] | Reference |
| Primary | 0.91 (0.239) | 25.9% [24.4-27.4] | -1.5% (p=0.242) |
| Junior high | 0.74 (0.001) | 22.6% [21.0-24.2] | -4.7% (p<0.001) |
| Senior high | 0.64 (<0.001) | 20.4% [17.9-22.8] | -7.0% (p<0.001) |
| Technical | 0.37 (0.051) | 13.6% [3.1-24.1] | -13.7% (p=0.012) |
| Middle technical | 0.54 (0.001) | 18.3% [14.1-22.4] | -9.1% (p<0.001) |
| Senior technical | 0.70 (0.049) | 21.8% [17.1-26.5] | -5.6% (p=0.040) |
| University or higher | 0.56 (0.012) | 18.7% [13.1-24.3] | -8.7% (p=0.006) |
| Income per capita (Yuan) |  |  |  |
| 0 - 4,747 | Reference | 29.6% [26.2-33.0] | Reference |
| 4,748 - 10,887 | 0.80 (0.035) | 25.7% [24.1-27.4] | -3.8% (p=0.039) |
| 10,888 - 17,631 | 0.71 (0.002) | 23.8% [22.0-25.6] | -5.8% (p=0.003) |
| 17,632 - 26,937 | 0.64 (<0.001) | 22.2% [20.2-24.1] | -7.4% (p<0.001) |
| 26,938 - 50,968 | 0.60 (<0.001) | 21.3% [18.7-23.8] | -8.3% (p<0.001) |
| >50,968 | 0.49 (0.011) | 18.4% [11.7-25.2] | -11.1% (p<0.001) |
| Age |  |  |  |
| Change/10 years of age | 0.91 (0.002) | Trend shown in Fig 2 | -1.4% (p=0.002) |
| Exercise |  |  |  |
| Never | Reference | 25.6% [24.4-26.9] | Reference |
| Up to 1 session/week | 0.86 (0.406) | 23.4% [18.4-28.4] | -2.2% (p=0.395) |
| 1-2 sessions/week | 0.82 (0.147) | 22.6% [18.7-26.5] | -3.1% (p=0.136) |
| 3-5 sessions/week | 0.79 (0.022) | 21.9% [19.2-24.8] | -3.7% (p=0.019) |
| ≥6 sessions/week | 0.80 (0.001) | 22.2% [20.6-23.8] | -3.5% (p=0.001) |
| Smoking (number of cigarettes) |  |  |  |
| None | Reference | 23.7% [22.7-24.6] | Reference |
| 1-10 per day | 0.87 (0.406) | 22.8% [20.0-25.7] | -0.1% (p=0.570) |
| 11-20 per day | 0.82 (0.147) | 25.7% [23.2-28.3] | 2.0% (p=0.133) |
| 21-30 per day | 0.79 (0.022) | 31.9% [25.5-38.4] | 8.2% (p=0.013) |
| >30 per day | 0.80 (0.001) | 31.3% [24.8-37.9] | 7.6% (p=0.024) |
| Regions |  |  |  |
| 1. Xicheng, Beijing | Reference | 8.9% [5.4-12.3] | Reference |
| 2. Yunhe, Cangzhou | 1.16 (0.631) | 10.1% [6.6-13.5] | 1.2 (0.630) |
| 3. Runzhou, Zhenjiang | 1.17 (0.636) | 10.1% [6.0-14.2] | 1.3 (0.638) |
| 4. Changning, Shanghai | 1.31 (0.363) | 11.2% [7.2-15.3] | 2.3 (0.365) |
| 5. Keqiao, Shaoxing | 2.77 (<0.001) | 20.5% [16.3-24.7] | 11.7 (<0.001) |
| 6. Meilie, Sanming | 3.62 (<0.001) | 25.0% [19.1-30.9] | 16.1 (<0.001) |
| 7. Yinzhou, Tieling | 5.46 (<0.001) | 32.9% [25.9-39.9] | 24.0 (<0.001) |
| 8. Lanshan, Linyi | 5.67 (<0.001) | 33.7% [26.3-40.9] | 24.8 (<0.001) |
| 9. Wuhou, Chengdu | 1.71 (0.065) | 14.0% [9.5-18.6] | 5.2 (0.067) |
| 10. Hongshan, Chifeng | 3.54 (<0.001) | 24.6% [20.0-29.2] | 15.7 (<0.001) |
| 11. Wudang, Guiyang | 4.16 (<0.001) | 27.5% [21.0-34.0] | 18.7 (<0.001) |
| 12. Chengbei, Xining | 5.28 (<0.001) | 32.2% [25.3-39.1] | 23.3 (<0.001) |
| 13. Yijiang, Wuhu | 1.65 (0.096) | 13.6% [9.3-17.8] | 4.7 (0.095) |
| 14. Yuhu, Shaoshan | 3.18 (<0.001) | 22.7% [17.9-27.5] | 13.8 (<0.001) |
| 15. Yiling, Yichang | 4.23 (<0.001) | 27.8% [21.8-33.8] | 19.0 (<0.001) |
| 16. Qing Yunpu, Nanchang | 4.87 (<0.001) | 30.5% [24.7-36.3] | 21.7 (<0.001) |
| 17. Changyi, Jilin | 5.73 (<0.001) | 33.9% [27.4-40.4] | 25.1 (<0.001) |
| 18. Pudong, Shanghai | 1.49 (0.152) | 12.5% [9.0-16.0] | 3.7 (0.143) |
| 19. Shengzhou, Shaoxing | 1.62 (0.085) | 13.4% [9.5-17.3] | 4.6 (0.079) |
| 20. uanghua, Cangzhou | 2.11 (0.011) | 16.6% [11.9-21.4] | 7.8 (0.010) |
| 21. Jurong, Zhenjiang | 2.43 (0.002) | 18.6% [13.5-23.7] | 9.7 (0.002) |
| 22. Shaxian, Sanming | 3.60 (<0.001) | 24.9% [19.2-30.5] | 16.0 (<0.001) |
| 23. Pinggu, Beijing | 4.37 (<0.001) | 28.4% [23.3-33.6] | 19.6 (<0.001) |
| 24. Xifeng, Tieling | 7.70 (<0.001) | 40.3% [33.3-47.3] | 31.5 (<0.001) |
| 25. Yinan, Linyi | 9.11 (<0.001) | 44.2% [36.9-51.4] | 35.3 (<0.001) |
| 26. Xinjin, Chengdu | 1.35 (0.354) | 11.5% [7.1-15.9] | 2.6 (0.359) |
| 27. Kaiyang, Guiyang | 6.33 (<0.001) | 36.0% [29.5-42.5] | 27.1 (<0.001) |
| 28. Huangzhong, Xining | 7.22 (<0.001) | 38.9% [31.8-45.9] | 30.0 (<0.001) |
| 29. Keshiketeng, Chifeng | 13.13 (<0.001) | 52.7% [46.6-58.8] | 43.8 (<0.001) |
| 30. Yidu, Yichang | 2.36 (0.003) | 18.1% [13.4-22.9] | 9.3 (0.002) |
| 31. Fanchang, Wuhu | 2.41 (0.002) | 18.5% [13.7-23.3] | 9.6 (0.002) |
| 32. Panshi, Jilin | 3.90 (<0.001) | 26.3% [19.6-33.1] | 17.5 (<0.001) |
| 33. Xiangtan, Shaoshan | 5.39 (<0.001) | 32.6% [27.4-37.8] | 23.7 (<0.001) |
| 34. Xinjian, Nanchang | 5.92 (<0.001) | 34.6% [25.2-43.9] | 25.7 (<0.001) |

<sup>a</sup> 95% confidence intervals.

#### The average effect of *Management* was estimated from the Main Effects logistic regression model of *Uncontrolled Hypertension*

Given the strong emphasis of the management program on the importance of regular compliance, at least some of the effect of management was expected to be due to improved compliance among managed patients. The results show that the effect of management was independent of compliance in the surveyed population, with no significant interaction (Table 4) or mediation effects (Table 5). To test for mediation, the compliance term was removed from the main effects regression model. The impact of management was not changed substantively in the absence of the compliance term (Table 5). Management was associated with a reduction of 8.6 percentage points (95% CI: −11.1 to −6.1) in uncontrolled hypertension if compliance was included in the model; without adjustment for compliance the reduction was 9.6 percentage points (95%CI: −12.1 to −7.0).

**Table 5.** Test for mediation in the impact of Management through Compliance.

| Type of estimate | Predicted estimates [95% CI] |
| --- | --- |
| Management odds ratio |  |
| • In a model with *Compliance* | 0.59 [0.51 to 0.68] |
| • In a model without *Compliance* | 0.57 [0.49 to 0.65] |
| Predicted failed control |  |
| • In a model with *Compliance* |  |
| Not managed | 31.0% [28.7 to 33.3] |
| Managed | 22.4% [34.1 to 47.4] |
| • In a model without *Compliance* |  |
| Not managed | 31.8% [29.5 to 34.0] |
| Managed | 22.2% [21.2 to 23.2] |
| Effect of management (%) |  |
| • In a model with *Compliance* | -8.6% [-11.1 to -6.1] |
| • In a model without *Compliance* | -9.6% [-12.1 to -7.0] |

#### The impact of management was compared in two Main Effects models: in one model *Compliance* was included as an independent variable, while in the second model the *Compliance* term was excluded

The effects of most other independent variables were in line with expectations (Table 4). Predictors of improved hypertension control included compliance, wellbeing, education, income, age and three or more exercise sessions per week, while smoking over 20 cigarettes per day was associated with reduced control. Predicted trends for wellbeing and age are shown in Fig 2 and S2 Fig. Levels of disease control also varied among regions (Table 4), with the most favorable outcomes predicted for Xicheng (8.9% uncontrolled hypertension), Yunhe (10.1%), Runzhou (10.1%), Changning (11.2%), Wuhou (14.0%), Yijiang (13.6%), Pudong (12.5%), Shengzhou (13.4%) and Xinjin (11.5%). The highest levels of uncontrolled hypertension were in Keshiketeng (52.7%) and Yinan (44.2%).

The significant inverse association between uncontrolled hypertension and age points to poor hypertension control in younger people. The trend is consistent with previous studies in both China and the USA, which showed that younger adults were less likely to be treated for hypertension than older adults. The residual effect may be due to a more general failure among younger people to seek medical care, including treatment for comorbidities, which may cause hypertension, such as diabetes and chronic kidney disease [19,20] (see Fig 2-1 and Fig 2-2).

**Fig 2-1.**
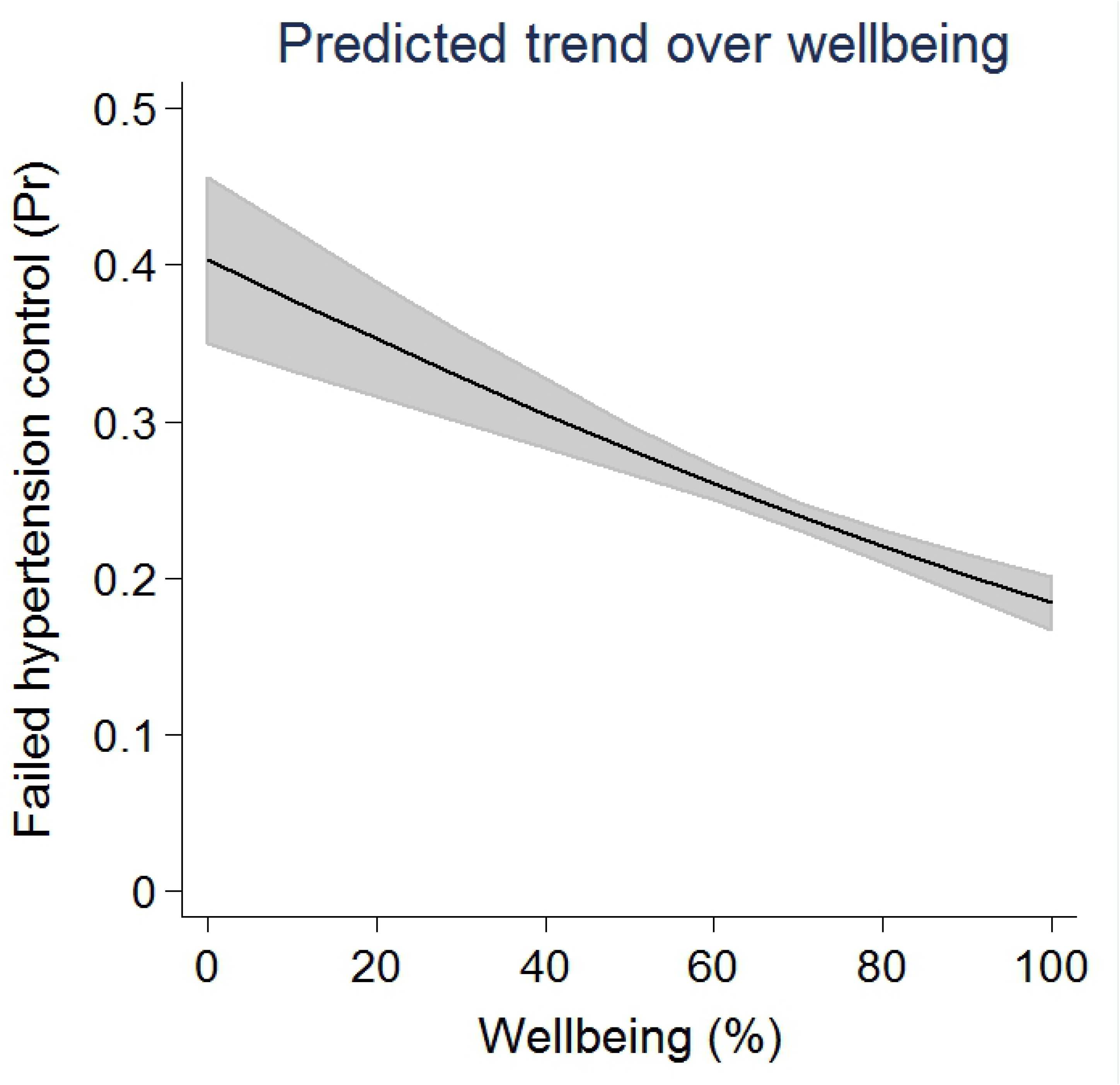
Predicted trends over wellbeing in failure to control hypertension.

**Fig 2-2.**
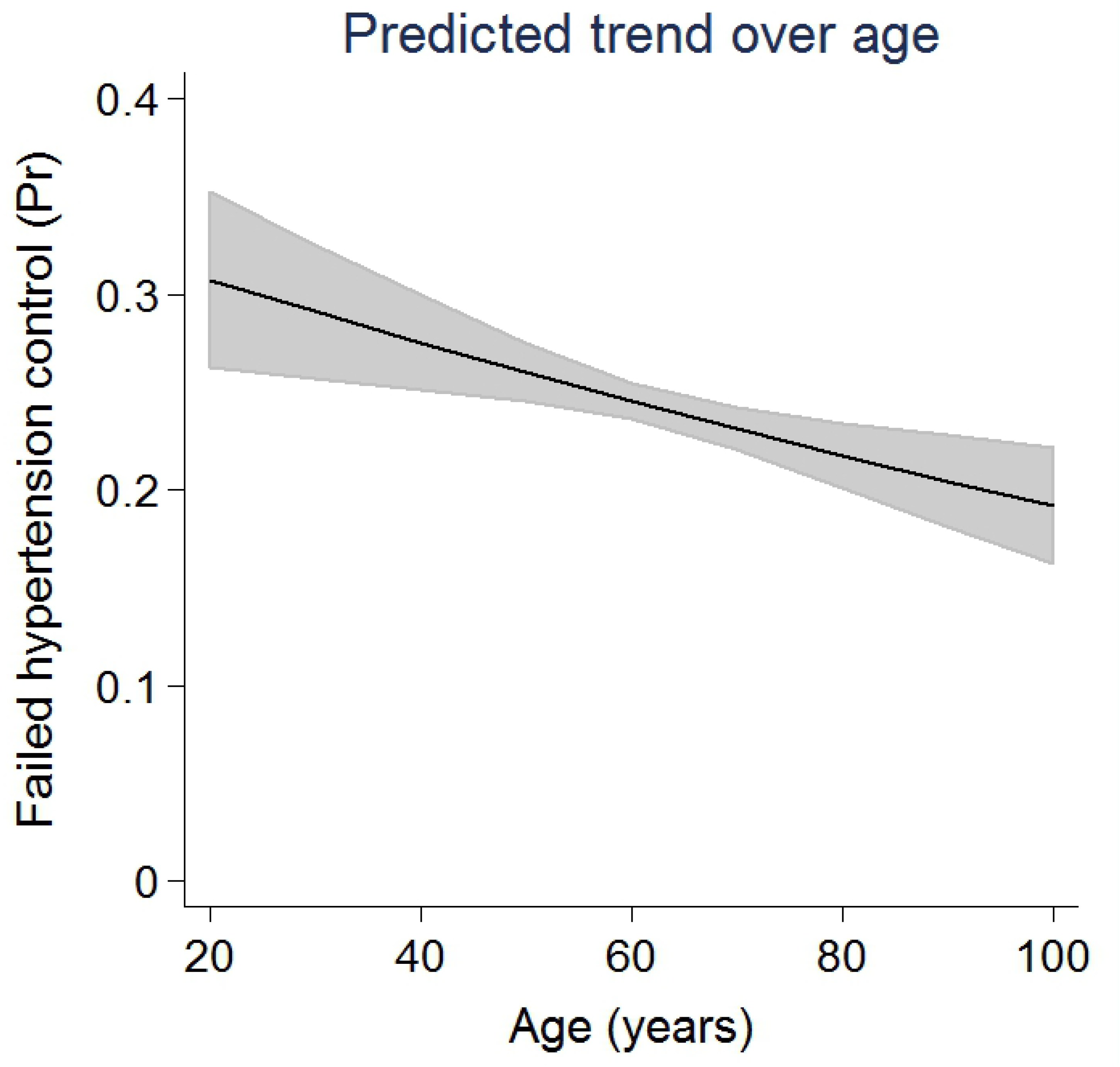
Predicted trends over age in failure to control hypertension. Age and wellbeing trends estimated using the Main Effects logistic regression model. Shaded areas represent 95% confidence intervals.

### Regional impact of management

The significant interaction between management and regions indicates that the benefit of the management program is not the same in all areas (χ^2^ [33] = 78.2, p<0.001, Table 3). Estimates for regional management effects are shown in Fig 3 and S3 Table. In sixteen regions, there were significantly lower rates of uncontrolled hypertension among managed patients, while in eighteen regions, no difference was detected between management groups. The strongest impact of management was observed in Wuhou and Shengzhou, with respective rate ratios of 0.19 and 0.20 (Fig 3 and S3 Table). Predicted difference proportions of *Uncontrolled Hypertension* between *Management* groups were derived from the Interaction Effects regression model (Table 2). The effect of *Management* in each location is indicated by black circles; vertical bars represent 95% confidence intervals. Estimates tagged with open circles mark those regions where no IMS systems have been established. Management rate ratio estimates for each region are listed in S3 Table and regional availability of IMS is shown in S4 Table.

**Fig 3.**
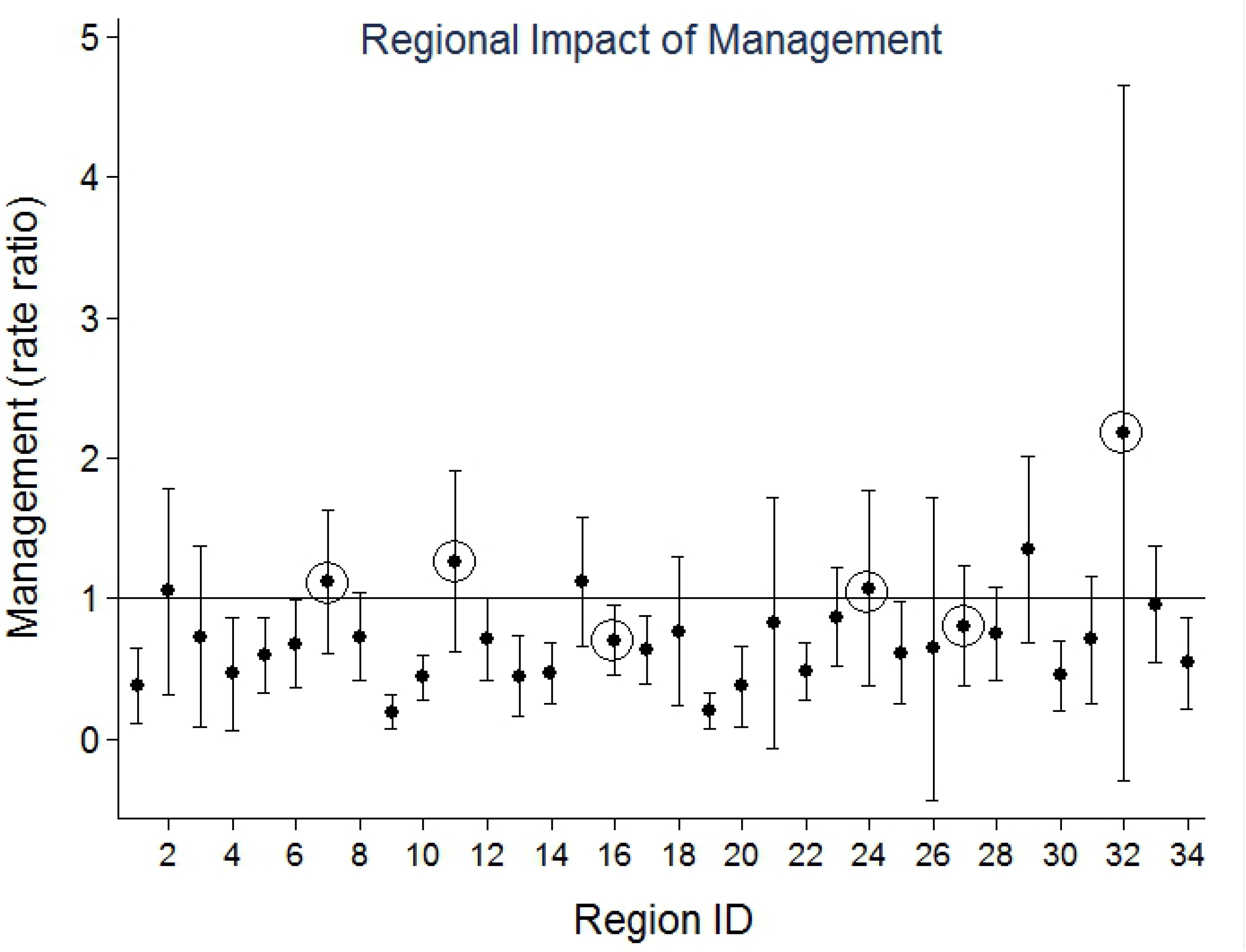
Marginal differences in *Management* across geographical regions. Predicted difference proportions of *Uncontrolled Hypertension* between *Management* groups were derived from the Interaction Effects regression model (Table 2). The effect of *Management* in each location is indicated by black circles; vertical bars represent 95% confidence intervals. Estimates tagged with open circles mark those regions where no IMS systems have been established. Management rate ratio estimates for each region are listed in S3 Table and regional availability of IMS is shown in S4 Table.

### Establishment of information management systems may contribute to regional differences in the effectiveness of the management program

Two important factors may account for the variability in regional management effects. One is the presence of an information management system (IMS), which is important according to the specification of hypertension management. We explored the possible link between regional variability in the impact of the management program and the presence of an established IMS. Regions with an established IMS are listed in S4 Table. Fig 3 shows that management rate ratios for these regions tended to cluster near the value of 1.00, which is the assumption of no effect under the null hypothesis. Management had no discernible impact on hypertension control in regions with no IMS (RR=0.93; 95% CI: 0.76-1.12), while in regions with established IMS, management had a strong impact (0.68, 95%CI: 0.62-0.75). Established IMS accounted for approximately 10% of the variability in regional management effects, as the change in deviance for the interaction term (IMS × management) represented 10% of the change in deviance for (regions × management) interaction terms (χ^2^=78.2, p<0.001).

The other possible factor is government funding, which has been well-reported in the literature in China [21–23]. However, in this study, no correlation was detected between regional management effects and government healthcare funding (S5 Fig). Related reported data on government funding for primary care and for total healthcare [22] is provided in S4 Table.

## 4. Discussion

This study examined the relationship between hypertension control and disease management under the PHSE program from care utilizers’ perspectives and found that providing management of hypertension significantly improved hypertension control.

Much of the literature on control of hypertension gives prominent roles to socio-economic factors, such as education, income and insurance, while taking healthcare accessibility and availability into account [6,23]. In this study, the above factors were not the main ones for hypertension control. Multi-regression analysis indicates that neither insurance scheme nor Hukou (which denotes urban and rural character of the surveyed hypertensives) were main factors for hypertension control. Such a result may imply that the EPHS program had played a role in equalization between urban and rural as well as that among different insurance schemes, which was like previous studies [24].

This study reveals major differences in the effectiveness of hypertension control among regions in China, which could guide policy makers and implementers in identifying where the health system needs strengthening. There are a number of possible reasons for these regional differences. According to the disease management specifications of hypertension from the PHSE issued by the National Health Commission (previously known as the National Health and Family Planning Commission) [25,26], one likely explanation is the efficiency of the integration of the management program with other medical services. Such integration is important in the treatment of patients who do not respond to medication. Appropriate referral for more intensive testing and treatment in a hospital or a specialist clinic is particularly important in the treatment of more severe hypertension or secondary hypertension, caused by diabetes, kidney disease or heart disease [20]. The ability to coordinate referrals as well as patient follow-up requires the establishment of effective information management systems (IMS) and government funding of the primary care system. This study has shown that the presence of an information management systems is a significant factor explaining regional differences in hypertension control. If some of the impact of management were mediated through IMS, it would be through providing improved access to advanced treatment for patients with severe or secondary hypertension. It follows that an effective system of referral can only be part of the solution and that more intensive treatment for severe or secondary hypertension must also be made available. It is possible that regional differences in the availability of more intensive treatment for patients with uncontrolled hypertension could account for some of the remaining differences in regional management effects.

Another possible reason for the variation of impact in different regions may be government funding. From international experience, investment by the government in an essential healthcare package is an important measure to address market failure in achieving the goal of universal health care [7,27]. In this study, a significant relationship between government funding and hypertension control was not detected. However, many studies in China discuss the impact of government funding on the PHSE program. In 2011, Chen Li et al. demonstrated that for the PHSE program, the financial investment could not match local responsibilities and roles of the central and local government were not clearly defined in funding and expenditure [20]. Such differences guide the need to further detect the regional differences of government investment, financial systems and expenditure on public health equalization programs at the primary care facilities level [28].

The failure to detect mediation effects could be explained by similarly high levels of compliance in both managed and unmanaged patients (85.4% and 78.1% respectively, Table 2). With high compliance rates among unmanaged patients, there was little opportunity for further improvement. In a different population with lower compliance rates outside the management program, we expect the effect of management to be more pronounced than estimated in this study because it would also include the contribution of improved medication use.

Younger people in the current study were also less likely to persist with medical treatment: they were less likely to participate in the management program and to comply with medication than older people. The finding of an inverse relationship between uncontrolled hypertension and age is not new [17,18]. It is noted that the inverse association between uncontrolled hypertension and age was detected in the regression model despite adjustment for management and compliance, which is consistent with the 2017 study by Li et al. [29]. Some younger people are part of a floating population and tend not to consider hypertension as a risk [30,31] which may contribute to poorer control among younger people. In China, the “floating population” of 250 million people mainly consists of young people and they suffer the highest illiteracy in China. As well, the health insurance system in China system is based at the local government level, and the floating population derives little benefit from it [31, 32].

As the survey was conducted by interviews, answers from respondents of hypertension management and control may not be as accurate as diagnosis by a health professional, although definitions such as hypertension control followed WHO guidelines. The findings of low management and control rates for young patients requires additional research, taking other social factors into account.

## 5. Conclusions

The PHSE program delivered through community health organizations has played a significant role in improving the control of hypertension and improved service delivery equalization on hypertension management between urban and rural and among different social-economic groups in China since its introduction in 2009. Based on a previous study [14], it is estimated that 7.31 million more patients received hypertension control from the PHSE in 2013. Patient outcomes have been improved by the program, independent of the degree of medication compliance, and demographic and socioeconomic differences, which reflected that equalization of service delivery was demonstrated. But the impact of the program on the control of hypertension varies across different regions in China and this is detected to be linked to the presence of a functioning health information system independent of the level of investment in health by the central government. This suggests that functioning health information systems be established in regions where they are not currently present and established systems be further strengthened to enhance the outcomes from the PHSE program. Lastly, implementation of the PHSE program needs to address the requirements of younger patients to improve the management of hypertension control to achieve higher levels of control.

## Author’s contribution

YZ conducted the initial logistic analysis and drafted the manuscript. MF undertook further statistical analysis and wrote the methodology and results section. JQ conceived and designed the study. KS contributed to the discussion and content of the paper. LZ, LM and CL participated in the writing of the manuscript. All authors read and approved the final manuscript.

## Acknowledgements

Margarita Kumnick from the Victoria Institute of Strategic Economics Studies provided professional editing of the whole paper. Prof. Meng Qingyue and Dr. Xu Jin, both from Peking University, gave the authors expert guidance in revising the paper.

## Supporting information

**S1 Table. Coding of questionnaires and questions (variables) from the survey questionnaire for this paper.**

**S2 Fig. Observed trends with age for compliance with medication as well as participation in the management program.**

**S3 Table. Variability of management effects over geographical localities.**

**S4 Table. Regional healthcare resources.**

**S5 Fig. Comparison of regional *Management* effects with total government healthcare spending.**

## References

1. Bloom G, Xingyuan G. Health sector reform: lessons from China. Soc Sc Med. 1997;45(3):351–60. doi: 10.1016/S0277-9536(96)00350-4

2. Yang G, Kong L, Zhao W, Wan X, Zhai Y, Chen LC, et al. Emergence of chronic non-communicable diseases in China. Lancet. 2008;372(9650):1697–705. doi: 10.1016/S0140-6736(08)61366-5

3. Yang G, Wang Y, Zeng Y, Gao G F, Liang X, Zhou M, et al. Rapid health transition in China, 1990-2010: findings from the Global Burden of Disease Study 2010. Lancet. 2013;381(9882):1987–2015. doi: 10.1016/S0140-6736(13)61097-1

4. Sinclair JAC. China’s health care reform. China Bus Rev. 1 Jul 2009. Available from: https://www.chinabusinessreview.com/chinas-healthcare-reform/

5. Yip WC, Hsiao WC, Chen W, Hu S, Ma J, Maynard A. Early appraisal of the huge and complex healthcare reform of China. Lancet. 2012;379(9818):833–42. doi:https://doi.org/10.1016/S0140-6736(11)61880-1

6. World Bank. Toward a healthy and harmonious life in China: stemming the rising tide of non-communicable diseases. Washington DC: 2011. Available from: http://www.worldbank.org/content/dam/Worldbank/document/NCD_report_en.pdf

7. World Health Organization. Package of essential noncommunicable disease interventions for primary health care in low-resource settings. Geneva: 2010. Available from: http://www.who.int/nmh/publications/essential_ncd_interventions_lr_settings.pdf

8. Department of Primary Care, National Health and Family Planning Commission of China. Policy explanation of how to implement essential health care package for public health equalization program in China. Beijing: 5 Sep 2017 [cited 2018 Sep 11]. In Chinese. Available from: http://www.nhfpc.gov.cn/jws/s3577/201709/9cb26d17f691491bb3bb0f139bbf3ff1.shtml

9. Xiao N, Long Q, Tang X, Tang S. A community-based approach to non-communicable chronic disease management within a context of advancing universal health coverage in China: progress and challenges. BMC Public Health. 2014;14 (Suppl 2):S2. doi: 10.1186/1471-2458-14-S2-S2

10. China National Health and Family Planning Commission. China health and family planning statistics yearbook 2014. Beijing: Peking Union Medical College Press; 2015.

11. Guo J, Zhu Y, Chen Y, Hu Y, Tang X, Zhang B. The dynamics of hypertension prevalence, awareness, treatment, control and associated factors in Chinese adults. J Hypertens. 2015;33(8):1688–96. doi: 10.1097/HJH.0000000000000594

12. Lu J, Lu Y, Wang X, Li X, Linderman GC, Wu C, et al. Prevalence, awareness, treatment, and control of hypertension in China: data from 1.7 million adults in population-based screening study (China PEACE Million Persons Project). Lancet. 2017;390(10112):2549–58. doi: 10.1016/S0140-6736(17)32478-9

13. Feng X, Pang M, Beard J. Health system strengthening and hypertension awareness, treatment and control: data from the China Health and Retirement Longitudinal Study. B World Health Organ. 2014;92:29–41. doi: 10.2471/BLT.13.124495

14. World Health Organization. The world health report 2008: primary health care: now more than ever. Geneva: World Health Organization; 2008 [cited 2018 Sep 11]. Available from: https://reliefweb.int/report/world/world-health-report-2008-primary-health-care-now-more-ever

15. Centre for Project Supervision and Management, National Health and Family Planning Commission, China. Annual report on essential public health services performance evaluation 2013. Beijing: 2014 [cited 2018 Mar 15]. Preprint. In Chinese.

16. Community Health Association of China. Report of periodical evaluation on the national essential public health programs 2015. Beijing: 2016 [cited 2018 Mar 15]. Preprint. In Chinese.

17. He J, Muntner P, Chen J, Roccella EJ, Streiffer RH, Whelton PK. Factors associated with hypertension control in the general population of the United States. Arch of Intern Med. 2002;162(9):1051–58. doi: 10.1001/archinte.162.9.1051.

18. Center for Health Statistics of the National Health and Family Planning Commission. An analysis report of the National Health Service 2013. Beijing: 2015 [cited 2018 Mar 15]. Available from: http://www.nhfpc.gov.cn/ewebeditor/uploadfile/2016/10/20161026163512679.pdf

19. Nwankuo T, Yoon SS, Burt V, Gu Q. Hypertension among adults in the United States: National Health and Nutrition Examination Survey, 2011-2012. NCHS Data Brief. 2013;Oct(133):1–8. doi: 10.1017/CBO9781107415324.004

20. Chen L, Shu Z, Yao L. Difficulties and strategies of the equality of essential public health services. China Health Econ. 2011;30(8):23–5. doi:10.03-10743(2011)08-0023-03

21. Zhou J, Meng Q, Miao Z. Study on the policy of the primary public health financing: based on the field survey in some disadvantaged rural areas. China Health Econ. 2011;30(6):7–9.

22. China National Health Development Research Centre, Department of Primary Care, National Health and Family Planning Commission of China. Monitoring and survey report of the 34 key contact pilot areas for primary care comprehensive reform of China. Beijing: 2015 [cited 2017 Jul 25]. Preprint. In Chinese.

23. Muntner P, Gu D, Wu X, Duan X, Wenqi G, Whelton PK, et al. Factors associated with hypertension awareness, treatment, and control in a representative sample of the Chinese population. Hypertension. 2004;43(3):57885. doi: 10.1161/01.HYP.0000116302.08484.14.

24. Hou Z, Meng Q, and Zhang Y. Hypertension Prevalence, Awareness, Treatment, and Control Following China’s Healthcare Reform. American Journal of Hypertension. 2016;29(4):428–431.

25. Ministry of Health, China; Ministry of Finance, China; State Administration of Traditional Chinese Medicine, China. Notice of implementing essential public health service equalization. Beijing: 2013 [cited 2017 Jul 25]. Available from: http://www.nhfpc.gov.cn/jws/s3577/201306/b035feee67f9444188e5123baef7d7bf.shtml

26. National Health and Family Planning Commission, China. National basic public health service delivery guideline of China. Beijing: 2013 [cited 2017 Jul 25]. Available from: http://www.moh.gov.cn/jws/s3577/201306/b035feee67f9444188e5123baef7d7bf.shtml

27. James MP. Government intervention in the markets for education and health care: how and why. NBER Working Paper No. 4916. Cambridge: 1995. Available from: http://www.nber.org/papers/w4916.pdf

28. Mendis S, Al Bashir I, Dissanayake L, Varghese C, Fadhil I, Marhe E, et al. Gaps in capacity in primary care in low-resource settings for implementation of essential noncommunicable disease interventions. Int J Hypertens. 2012;(584041). doi: 10.1155/2012/584041

29. Li Y, Zeng X, Liu J, Liu Y, Liu S, Yin P, et al. Can China achieve a one-third reduction in premature mortality from non-communicable diseases by 2030. BMC Medicine. 2017;15(1):132. doi: 10.1186/s12916-017-0894-5

30. Gooding HC, McGinty S, Richmond TK, Gillman MW, Field AE. Hypertension awareness and control among young adults in the National Longitudinal Study of Adolescent Health. J Gen Intern Med. 2014;29(8):1098–104. doi: 10.1007/s11606-014-2809-x

31. Zhu Y. China’s floating population and their settlement intention in the cities: beyond the Hukou reform. Habitat Int. 2007;31(1):65–76. doi: 10.1016/j.habitatint.2006.04.002

32. Armstrong TD. China’s “floating population”. South Calif Int Rev. 12 October 2013 [cited 2017 Aug 27]. Available at: http://scir.org/2013/10/chinas-floating-population/

